# Razor: annotation of signal peptides from toxins

**DOI:** 10.1101/2020.11.30.405613

**Authors:** Bikash K. Bhandari, Paul P. Gardner, Chun Shen Lim

## Abstract

**Motivation:** Signal peptides are responsible for protein transport and secretion and are ubiquitous to all forms of life. The annotation of signal peptides is important for understanding protein translocation and toxin secretion and evolution.

**Results:** Here we explore the features of these signal sequences from eukaryotic proteins. Strikingly, we find that the signal peptides from secretory toxins have common features across kingdoms, supporting the idea of horizontal gene transfer or convergence of toxin genes across kingdoms. We leverage these features to build Razor, a simple yet powerful tool specialised in identifying signal peptides from toxins using the first 23 N-terminal residues. We demonstrate the usability of Razor by analysing all the sequences reviewed by UniProt. Indeed, Razor is able to identify toxins using their N-terminal sequences only. Strikingly, we also discover that many defensive proteins across kingdoms harbour a toxin-like signal peptide; some of these defensive proteins have emerged through convergent evolution, e.g. defensin and defensin-like protein families, and phospholipase families. In sum, Razor uses an approach independent of homology search to identify novel and known toxin classes across species using N-terminal residues.

**Availability and implementation:** Razor is available as a web application (https://tisigner.com/razor) and a command-line tool (https://github.com/Gardner-BinfLab/Razor).

## Introduction

Secretory proteins are translocated in the secretory pathway with the assistance of a short peptide extension at the N-terminus. This special targeting peptide is known as the Signal Peptide (SP) (von Heijne, 1990). Secretory pathways and their corresponding SPs have evolved across organisms to carry out different functions (Hegde and Bernstein, 2006; Owji *et al*., 2018). Despite being ubiquitous across all domains of life, SPs do not share a consensus. Nevertheless, a SP usually consists of three regions: a positively charged domain (N-region), a hydrophobic core (H-region), followed by a polar but electrically neutral domain (C-region) containing a cleavage site (von Heijne, 1985, 1990; Nielsen and Krogh, 1998). Apart from translocating proteins, SPs are also responsible for several other roles, such as in regulatory functions, antigen presentation, and some human diseases (Borrego *et al*., 1998; Datta*et al*., 2007; Owji*et al*., 2018).

An important group of secretory proteins is toxins, whose precursors almost always contain SPs (Fry *et al*., 2009). Toxins have evolved in all domains of life primarily as a defense mechanism or for predation (Casewell *et al*., 2013). Furthermore, several organisms in the animal kingdom have evolved to create venoms, which consist of a complex mixture of different types of toxins, usually with a specialised apparatus to facilitate their delivery. Such adaptations may have evolved through convergence or duplication and neofunctionalisation (Casewell, 2020). However, a recent study found that at least five toxin gene families were horizontally transferred from bacteria and fungi to centipedes (Undheim and Jenner, 2021), suggesting common features exist in these gene families. Besides, the pharmacological actions of toxins on living cells are often employed to develop anti-toxins, novel drugs, and pathogen-resistant transgenic crops (King, 2011; Estrada *et al*., 2007; Bidondo *et al*., 2019; Samy*et al*., 2017; Li*et al*., 2018). Hence, annotating SPs is essential in the functional and structural studies of proteins in fundamental research, commercial, and pharmaceutical industries. In addition, understanding the presence or absence of SPs in the genes of interest is critical for choosing the appropriate recombinant protein expression and purification systems, as the intracellular accumulation of secretory proteins and toxins may be toxic to the host cells. Indeed, the ability of SPs to translocate proteins has been utilised in recombinant protein expression systems for high quality and quantity results (Futatsumori-Sugai and Tsumoto, 2010; Cho *et al*., 2019; Karyolaimos *et al*., 2019; Peng *et al*., 2019).

Despite the immense use cases of toxins, there are very few tools to predict them, such as ClanTox, ToxinPred, TOXIFY, and ToxClassifier, some being specialised such as SpiderP for spider toxins (Naamati *et al*., 2010; Gupta *et al*., 2013; Wong *et al*., 2013; Gacesa *et al*., 2016; Cole and Brewer, 2019). Moreover, these methods are based on the properties of the mature peptides (or the propeptides), rather than the SPs. To address these issues, we first examined the features of SPs from eukaryotic proteins and toxins. We then exploited those features to build Razor, a new tool for annotating SPs. We have optimised the command-line version of Razor for high-throughput analysis and used it to predict new SPs by scanning all the sequences reviewed by UniProt (UniProt Consortium, 2019). We were able to predict novel toxins and defensive proteins using only the first 23 N-terminal residues, as evidenced by the protein family annotations.

## MATERIALS AND METHODS

### Datasets

We retrieved the training dataset for the state-of-the-art SP prediction program SignalP 5.0, which is a curated set of the N-terminal sequences from all domains of life (Almagro Armenteros *et al*., 2019). To get the full sequences and annotations of eukaryotic proteins, we used UniProt ‘s ID mapping service (UniProt Consortium, 2019) and obtained 17,264 fully annotated sequences, of which 2,609 sequences have been experimentally validated to harbour functional SPs. These sequences were used to build a generic, eukaryotic SP classifier. For feature analysis, we clustered these sequences (60 N-terminal residues) at an identity threshold of 70% using CD-HIT v4.8.1 (Fu *et al*., 2012). A single representative sequence was retained for each cluster to reduce sequence redundancy (Supplementary Table S1).

To build a classifier specialised for annotating toxin SPs, we manually curated a separate positive set using the dataset from the animal toxin annotation project (Jungo *et al*., 2012) and a subset from the above training set. Other SPs were assigned as a negative set. We then clustered the sequences as above and analysed the representative sequences (Supplementary Table S1).

The SP classifiers were compared using an independent test set retrieved from UniProt on 16 February 2021. In particular, the eukaryotic SP classifier was evaluated using 241 SPs with experimental evidence and 52,055 non-SPs, whereas the toxin SP classifier was evaluated using a subset of this independent set (toxin SPs=47, non-SPs=52,055). We also scanned the reviewed sequences from UniProt (N=561,776, retrieved on 2 September 2020).

### Bit score

The bit scores of the N-terminal residues were computed as:

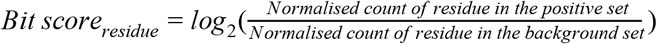

For eukaryotic proteins, the positive set and the background set were SPs and non-SPs, respectively. For toxins, the positive set and the background set were toxin SPs and non-toxin SPs, respectively.

### Protein sequence properties

The standard protein sequence properties, implemented in BioPython, were calculated using the Bio.SeqUtils.ProtParam module v1.73 (Cock *et al*., 2009). These features include GRand AVerage of hydropathicitY (GRAVY), Flexibility, Helix, Sheet and Turn propensities, Instability Index, Aromaticity, and Isoelectric Point. An additional feature included is the Solubility-Weighted Index (SWI; (Bhandari *et al*., 2020).

### SP classifiers

We built a random forest classifier based on several sequence features (GRAVY, flexibility, helix, and SWI), as well as the counts of residues (R, K, N, D, C, E, V, I, Y, F, W, L, Q, and P) of the first 30 N-terminal residues. The residues were chosen such that they maximised Matthew ‘s correlation coefficient (MCC) in five-fold cross-validations. After the cross-validation step, we generated five random forest models, which are used for scoring the N-terminal of a given sequence. The scores from these classifiers are comparable to the S-score of SignalP 4.0 except that our scores are non-position-specific (Petersen *et al*., 2011).

For the prediction of the cleavage site, we took a total of 30 residues such that the cleavage site is aligned in between positions 15 and 16 in order to capture the major differences in residue distribution around the cleavage site. We built a 20×30 matrix and populated it with the hydrophobicity scale (Kyte and Doolittle, 1982) as initial weights. We then used multi-objective simulated annealing (Kirkpatrick *et al*., 1987) at each position such that the new weights maximised the AUC and precision-recall curve based on the training set. The scoring of the cleavage site (C-score) is done using the random forest classifier trained on the aligned set encoded using the optimised weight matrix. Small limitation of our approach is that we are unable to detect the correct cleavage site if it is located before the 15th position. Yet, based on training data, this is rarely observed (N=13).

After detecting the cleavage site, the final score for classification (Y-score) is the geometric mean 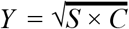 where S is the S-score and C is the max of C-scores along the sequence. For the final classifier, we chose a threshold of Y-score that maximised the MCC after five-fold cross-validations (MCC=0.914) on the training set.

We then built models specialised in annotating the toxin SPs based on hydrophobicity, SWI, flexibility, and turn. These features were selected such that they maximised the MCC using five-fold cross-validations on the training set. The N-terminal length of 23 was found to generate the maximum median MCC score for the toxin SP classifier (MCC=0.741, see also Supplementary Table S2). Similar to the SP prediction models, the toxin SP classifiers consist of five models each.

### Performance measures

We use MCC as a measure of performance to correctly identify eukaryotic SPs. We also use cleavage site precision (*CS*_*P*_ = *N* _*corr*_/*N*_*P*_) and recall (*CS*_*R*_ = *N* _*corr*_/*N*), where *N* _*corr*_is thenumber of the correctly identified cleavage site, *N*_*P*_is the number of predicted SPs and *N* is the number of SPs (Almagro Armenteros *et al*., 2019; Savojardo *et al*., 2018).

### Tool

We developed Razor for annotating SPs using the eukaryotic and toxin SP classifiers (Fig 1). Razor accepts either a nucleotide sequence or a protein sequence. Sequences with a length of lower than 30 residues are padded with Serine (Ser, S), because it shows equal enrichment across all datasets, in particular after the H-region (Fig 2). Razor is available both as a command-line tool (https://github.com/Gardner-BinfLab/Razor) and a web application (https://tisigner.com/razor). For the web application, predictions from five models are displayed as stars. The final score is the median of scores from five models and is displayed along with the region for SP. A plot of C-scores along the sequence is also displayed along with the annotation for the cleavage site. In addition, we integrated the Razor web application with our protein expression and solubility optimisation tools, TIsigner and SoDoPE, respectively (Bhandari *et al*., 2020, 2021). Our web tools assist users in annotating SPs and protein domains, and making the decisions from gene cloning to protein expression and purification.

**Fig 1.**
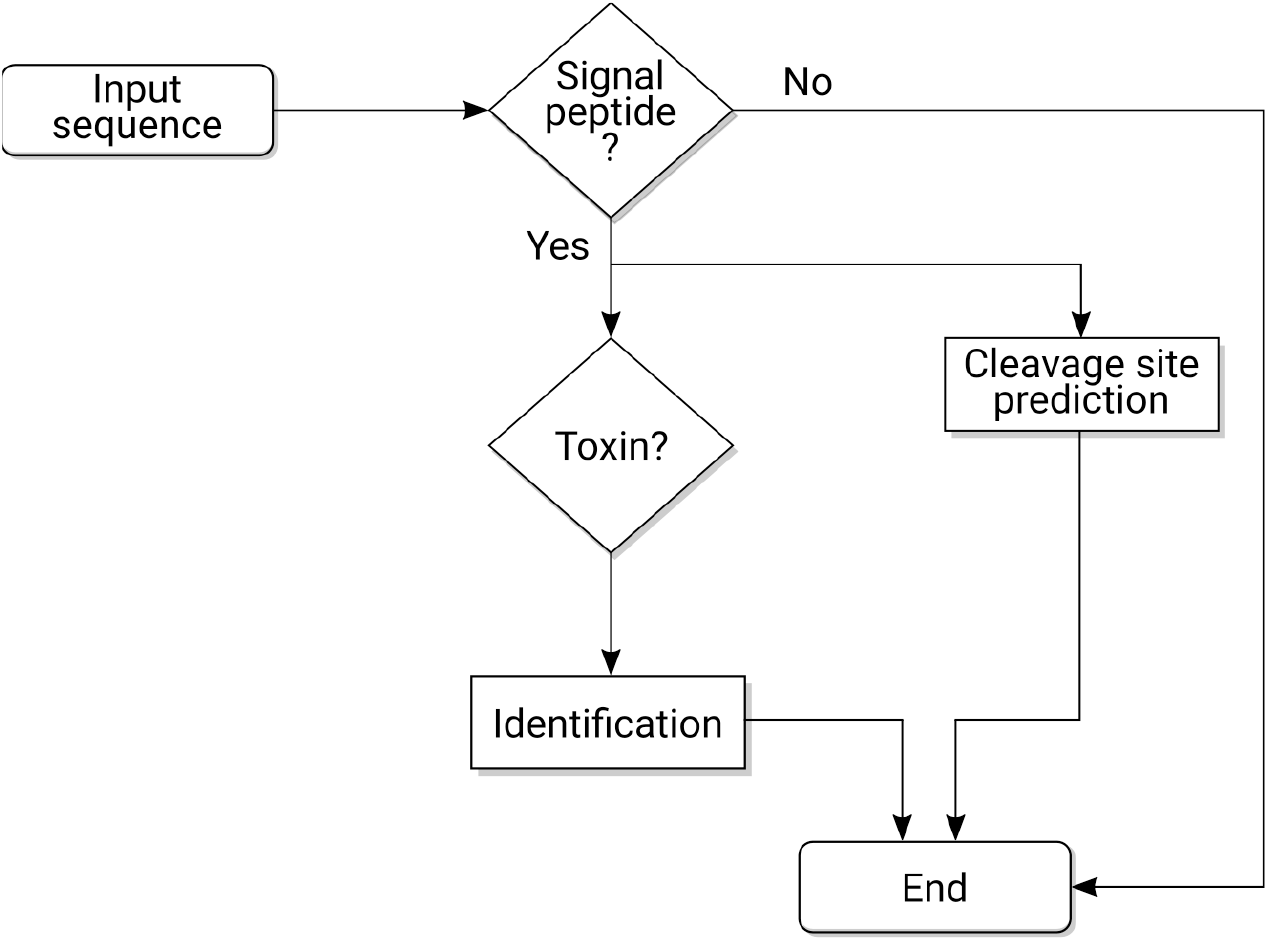
Flow chart of toxin SP classification using Razor.

**Fig 2.**
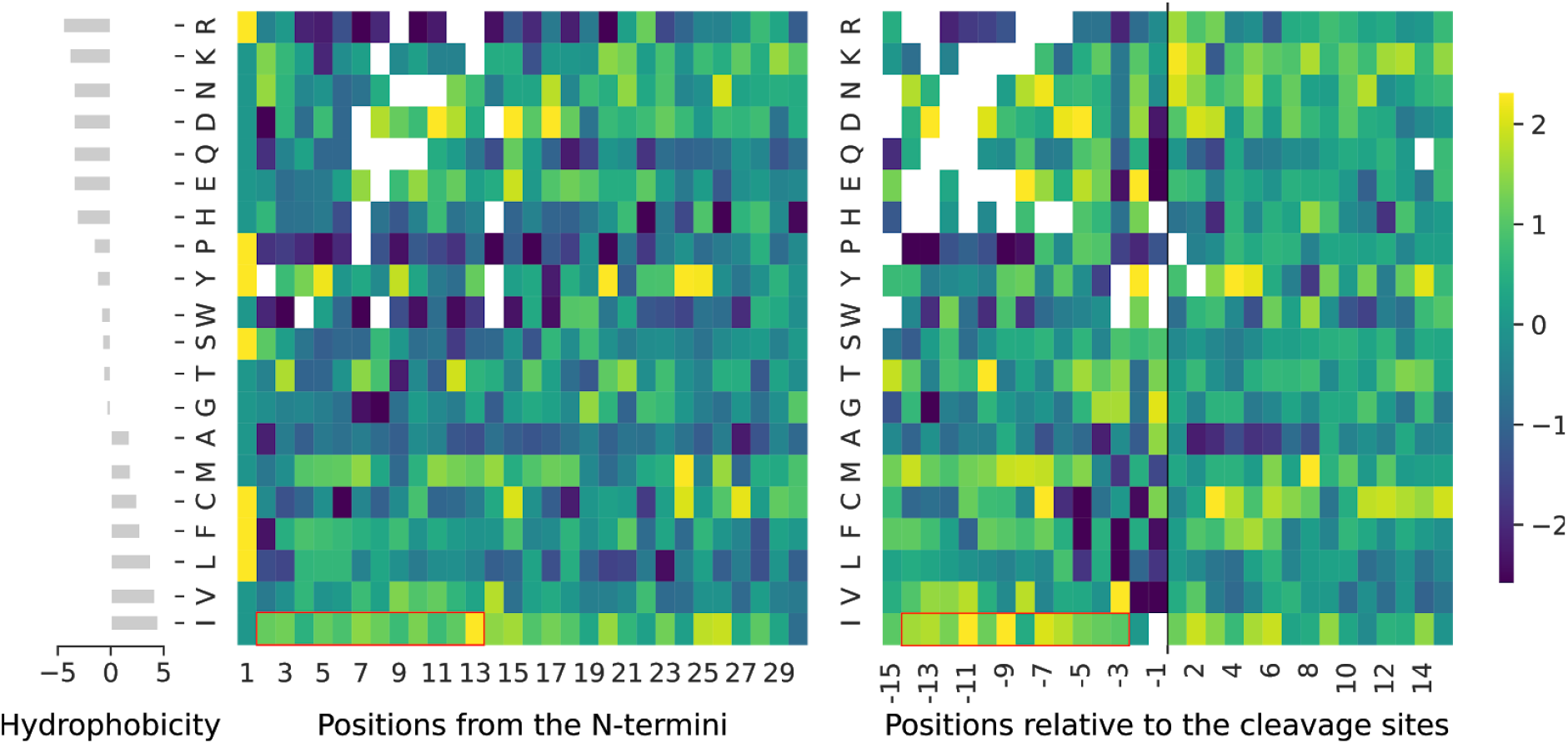
The Signal Peptides (SPs) from toxins are enriched with isoleucine residues in contrast to other eukaryotic SPs. The bar plot shows Kyte and Doolittle ‘s hydrophobicity scale. The heatmaps show the enrichment of residues in bit scores by aligning SPs from the N-termini (left) and at the cleavage sites (right, black vertical line). The unfilled, red rectangles indicate the enrichment of isoleucine residues (I). The white spaces correspond to the absence of residues at certain positions due to limited sample size (261 toxin SPs and 1,738 non-toxin SPs that have been experimentally validated).

### Statistical analysis

Data analysis was performed using pandas v1.0.3 (McKinney, 2010). Hydrophobicity and SWI were smoothed for the classifier training using the Savitzky-Golay filter implemented in SciPy v1.4.1 (Virtanen *et al*., 2020). Random forest classifier and MCC computation were done using scikit-learn v0.23.1 (Pedregosa *et al*., 2011). Plots were generated using Matplotlib v3.1.3 and Seaborn v0.10.0 (Hunter, 2007; Waskom *et al*., 2020).

### Code and data availability

Jupyter notebooks for reproducing our analyses are available at https://github.com/Gardner-BinfLab/Razor_paper_2021. The source code for Razor, our SP annotation server can be found at https://github.com/Gardner-BinfLab/TISIGNER-ReactJS.

## RESULTS

### Toxin SPs have distinct sequence properties

We investigated the sequence composition of SPs by first aligning the sequences from the N-terminal residue or by centering at the cleavage sites, followed by computing bit scores for each residue (Fig 2). These approaches provide sufficient leverage to enumerate the tripartite domains of SPs (N-, H-, and C-domains). In general, hydrophobic residues are enriched towards the N-termini (H-region), which are characteristic features of SPs (von Heijne, 1990) (Supplementary Fig S1). Strikingly, the SPs of toxins show a strong abundance of isoleucine (I) and lack leucine (L) and alanine (A) residues in contrast to other eukaryotic SPs (Fig 2). This is supported by an amino acid composition analysis of the N-terminal subsequences (Supplementary Fig S2). We also analysed other features of these N-terminal subsequences, including GRAVY, structural flexibility, helix, sheet and turn propensities, instability index, aromaticity, isoelectric point, and SWI. Interestingly, isoelectric point appears as a prominent feature of toxin SPs (Supplementary Fig S3).

The cleavage sites mark the end of SPs and the beginning of the mature region (or the propeptide), which is a unique feature of SPs (Fig 2). By aligning the sequences at the cleavage sites, we observed a clear emergence of (−3,−1) rule preceding the cleavage sites, i.e. a distinctive presence of small and charged residues such as alanine (A) and valine (V) (von Heijne, 1983).

### Razor accurately predicts toxin SPs

By taking these important features into account, we built SP classifiers to annotate eukaryotic and toxin SPs using random forest (Fig 1). Only SPs with experimental evidence were used for training. We compared these classifiers using an independent test set, where, the MCC, and the cleavage site precision and recall of Razor for eukaryotic SP prediction were 0.405, 0.136, 0.596, respectively (SPs vs non-SPs, see Supplementary Fig S4, Table S3 and S4). More importantly, Razor outperforms state-of-the-art in toxin SP prediction, achieving an MCC score of 0.611, and the cleavage site precision and recall of 0.355 and 0.831, respectively (toxin SPs vs non-SPs, see Fig 3, and Supplementary Table S5 and S6).

**Fig 3.**
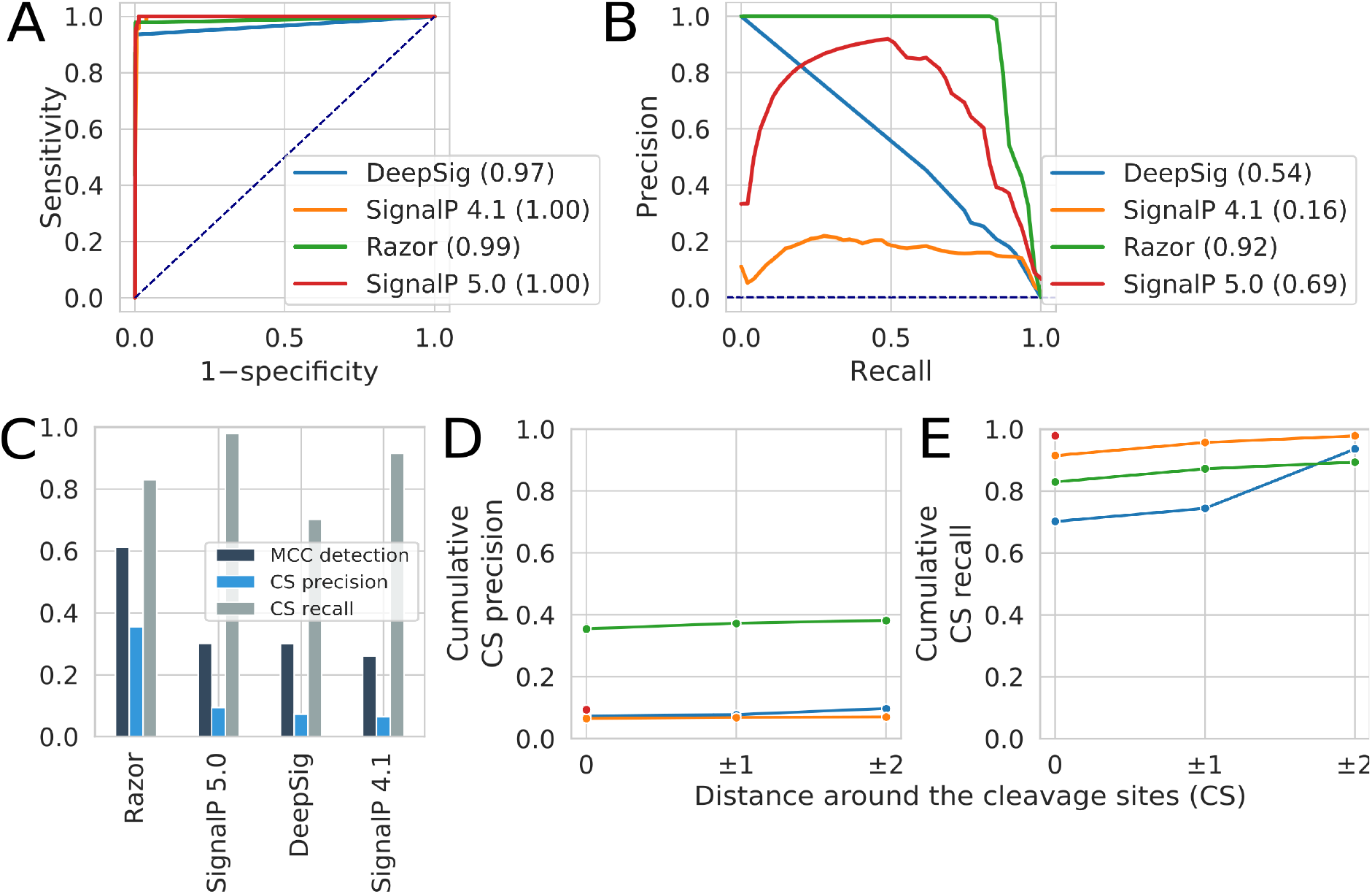
Razor outperforms other tools in predicting toxin SPs. Benchmarks were carried out using an independent test set (47 experimentally validated toxin SPs and 52,055 non-SPs). **(A)** Receiver operating characteristic curves **(B)** and precision recall curves **(C)** of the SP prediction tools. Areas under the curves are shown in parentheses. The dotted lines show the performance of a random classifier. **(C)** Matthew ‘s Correlation Coefficients (MCC) of the SP prediction tools. The cleavage site (CS) precisions **(D)** and recalls **(E)** of windows surrounding the cleavage sites are shown. Data are available in Supplementary Tables S5 and S6.

### Defensive proteins harbour a toxin-like SP

The training set for the toxin SP classifier was mainly composed of the SPs from animal toxins, e.g. snake three-finger toxins, scorpion toxins, and phospholipase A _2_, and plant toxins, e.g. ribosome-inactivating proteins (Fig 4A). To further assess our new toxin SP classifier, we scanned the reviewed sequences from UniProt (N=561,776). A total of 910 sequences were predicted positive from all SP detection models.

**Fig 4.**
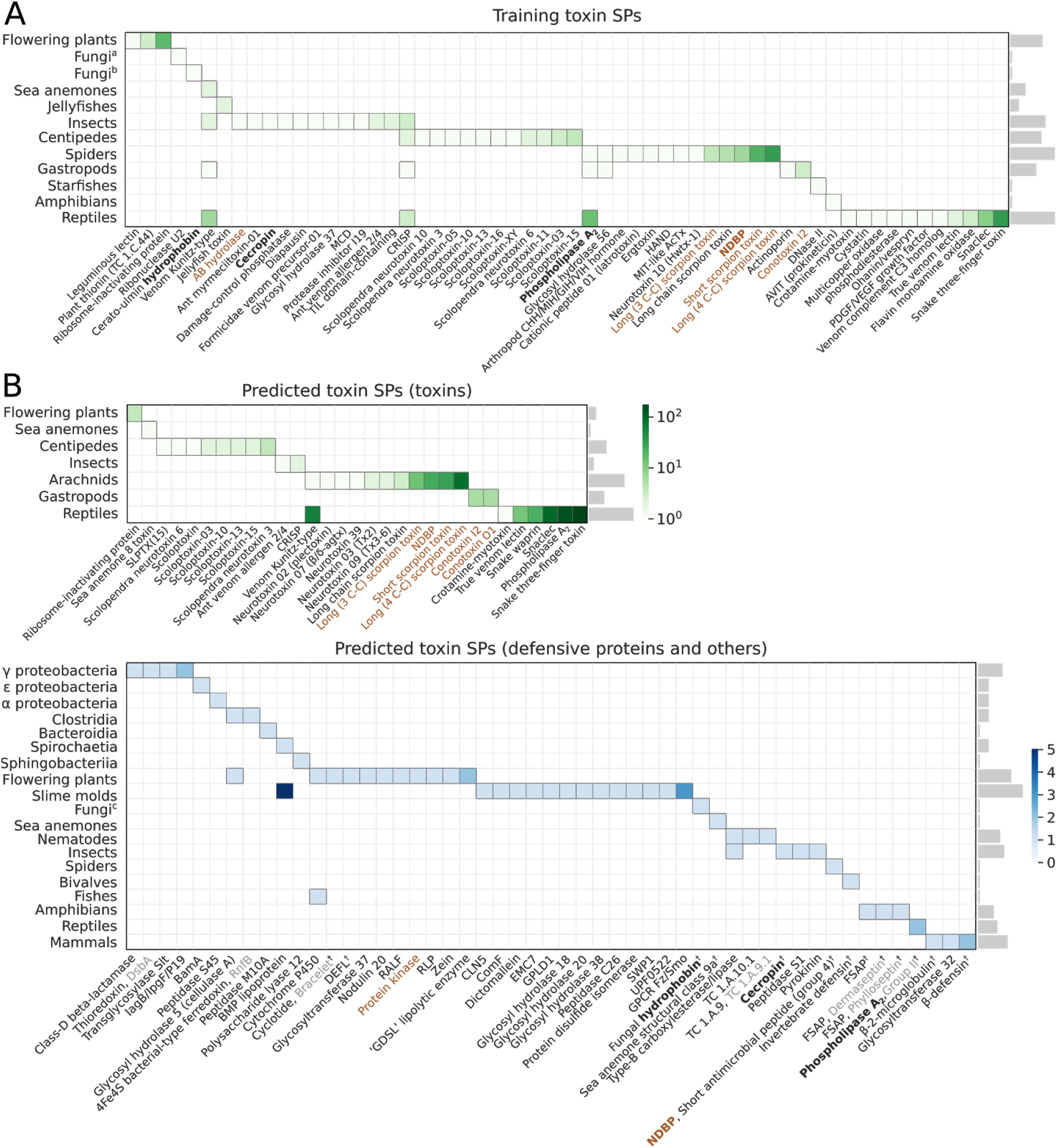
Razor identifies SPs from toxins along with several classes of defensive proteins. The reviewed sequences from UniProt were examined (N=561,776). **(A)** Heatmap shows the abundance of protein families in the training toxin sequences with SPs by taxa. A total of 237 of 261 training toxins had protein family annotations. **(B)** Heatmaps show the abundance of protein families in the sequences predicted to harbour toxin SPs. A total of 753 of 759 toxins predicted to harbour toxin SPs had protein family annotations (top). A total of 110 other types of proteins were predicted to harbour toxin SP, in which 76 of them had protein family annotations (bottom). The scale bars indicate the frequencies of protein families. Those protein families that have defensive properties are marked with † (bottom). Protein families that are in common between the training and predicted toxin SP sequences are bolded (bottom panel). Protein subfamily, family and superfamily are shown in grey, black and brown, respectively. Fungi^a^, Eurotiomycetes; Fungi^b^, Sordariomycetes; Fungi^c^, Agaricomycetes; CLN5, Ceroid-Lipofuscinosis Neuronal protein; ComF, Competence protein F; CRISP, Cysteine RIch Secretory Protein; DEFL, DEFensin Like; EMC7, ER membrane protein complex subunit 7; FSAP, Frog Skin Antimicrobial Peptide; GPLD1, Glycosyl-phosphatidylinositol-specific phospholipase D; HAND, Helical Arthropod-Neuropeptide-Derived; RALF, Rapid ALkalinization Factor; RLP, Receptor Like Protein; SLPTX, Scoloptoxin; UPF, Uncharacterised Protein Family.

In Fig 4, we excluded potential false positive hits, i.e. computationally annotated transmembrane proteins by UniProt (N=33). The remaining sequences were divided into two groups based on the presence or absence of toxin annotation. From these probable toxin SPs, 759 sequences had annotations for toxins. They included protein families such as scorpion toxin, phospholipase A_2_ and ribosome-inactivating protein (Fig 4B). The remaining 110 sequences had no annotations for toxins. These sequences were clustered at an identity threshold of 70%, which gave rise to 100 representative sequences. Interestingly, many of these proteins without toxin annotations have some defensive properties such as antibacterial peptides and cyclotides. Furthermore, other defensive proteins such as beta-defensin and defensin-like (DEFL) are the results of convergent evolution. For example, beta-defensin-like motifs are also found in toxins from lepidosauria (rattlesnakes and bearded dragons) and mammalia (platypus) (Fry *et al*., 2009, 2010; Whittington *et al*., 2008). This suggests why their SPs show some remote similarity with toxin SPs.

## DISCUSSION

We have studied the features of SPs from eukaryotic proteins. While SPs share a common hydrophobic nature, we have found several differences between toxin SPs and other eukaryotic SPs in their residue compositions and consequently the sequence properties. We have used these features to develop Razor for annotating eukaryotic SPs, which have specialised functionalities in annotating toxin SPs. Razor outperforms other sophisticated methods in predicting toxin SPs. Using Razor, we were able to predict several classes of probable toxins, which are yet to be annotated (Fig 4). Our predicted results consist of toxins and defensive proteins from diverse species, which gives us an overview of the source of toxins.

Since toxins and defensive proteins occur naturally in organisms to attack and neutralise foreign invaders, many of our predicted results include proteins involved in innate immune response and signalling. Some of the frequently observed biological processes of these proteins were ‘killing of cells of other organism [GO:0031640] ‘, ‘defense response to fungus [GO:0050832] ‘, ‘defense response to bacterium [GO:0042742] ‘ and ‘innate immune response [GO:0045087] ‘ (Supplementary Fig S5 and S6). Many toxins and defensive proteins are commercially important. For example, plant toxins such as defensin-like protein, animal toxins such as cecropin are used to develop disease-resistant transgenic crops (Stotz *et al*., 2009; Lacerda *et al*., 2014; Wu *et al*., 2016; Boccardo *et al*., 2019; Ali *et al*., 2018). Similarly, the cytotoxic activity of phospholipase A_2_ on cancer cells makes it a promising candidate for cancer therapy (Xiao *et al*., 2017; Hiu and Yap, 2020; Lomonte and Rangel, 2012).

Taken together, Razor uses an approach independent of homology search to identify known and novel toxin classes across species. Razor was able to identify previously unannotated SPs and a spectrum of toxins and defensive proteins simply using the first 23 N-terminal residues. This also suggests a possible evolutionary constraint on SPs driven by the specialisation of the toxin secretory systems (or convergent evolution), and supports the idea of horizontal gene transfer of several toxin gene classes (Undheim and Jenner, 2021). Therefore, accurate annotation of toxin SPs can enhance comparative genomics analysis and genome sequencing projects. Razor might also be useful in other research areas such as recombinant protein expression, toxicology, transgenics, and drug design.

## AUTHORS CONTRIBUTIONS

CSL conceived the study. BKB carried out the analysis, built Razor, and drafted the manuscript. CSL and PPG supervised the study. All authors reviewed, edited, and approved the manuscript.

## ACKNOWLEDGEMENTS

The authors thank Dr Astra Heywood for providing feedback on the figures and the Razor web application.

