## Supplementary Fig, Supplementary Table for "Razor: annotation of signal peptides from toxins"

**SUPPLEMENTARY MATERIALS**

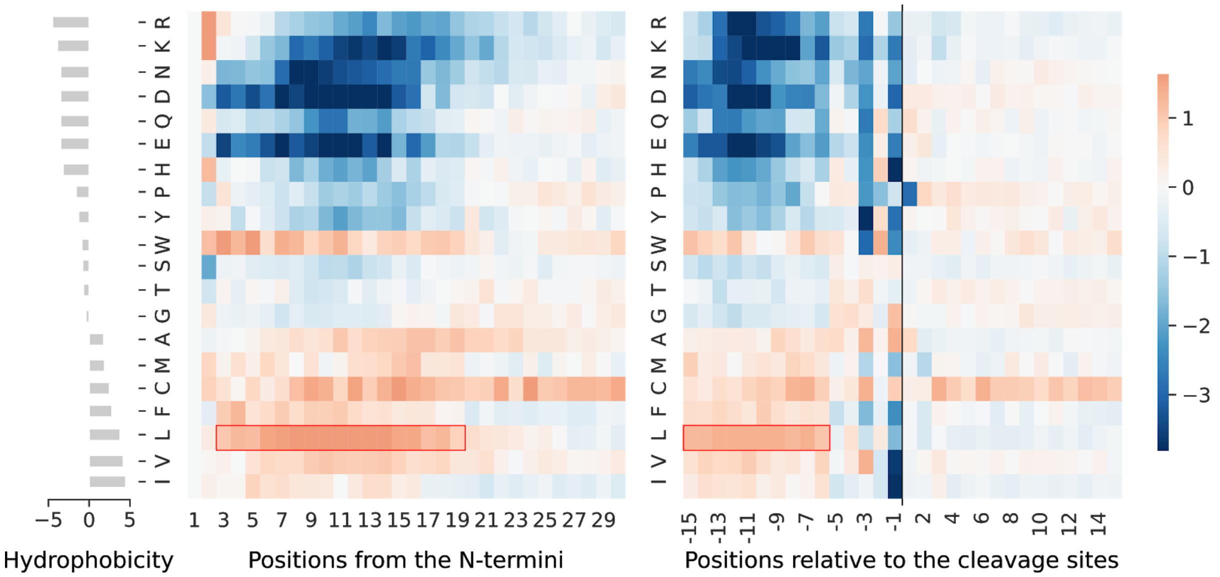


**Fig S1. Signal peptides (SPs) show a strong hydrophobic property (1,964 experimentally validated SPs, 13,237 non-SPs).** The bar plot shows Kyte and Doolittle's hydrophobicity scale. The heatmaps show the enrichment of residues in bit scores by aligning SPs from the N-termini (left) and at the cleavage sites (right, black vertical line). The (-3, -1) rule for the cleavage site motif is shown (left). The unfilled, red rectangles indicate the enrichment of leucine residues (L).

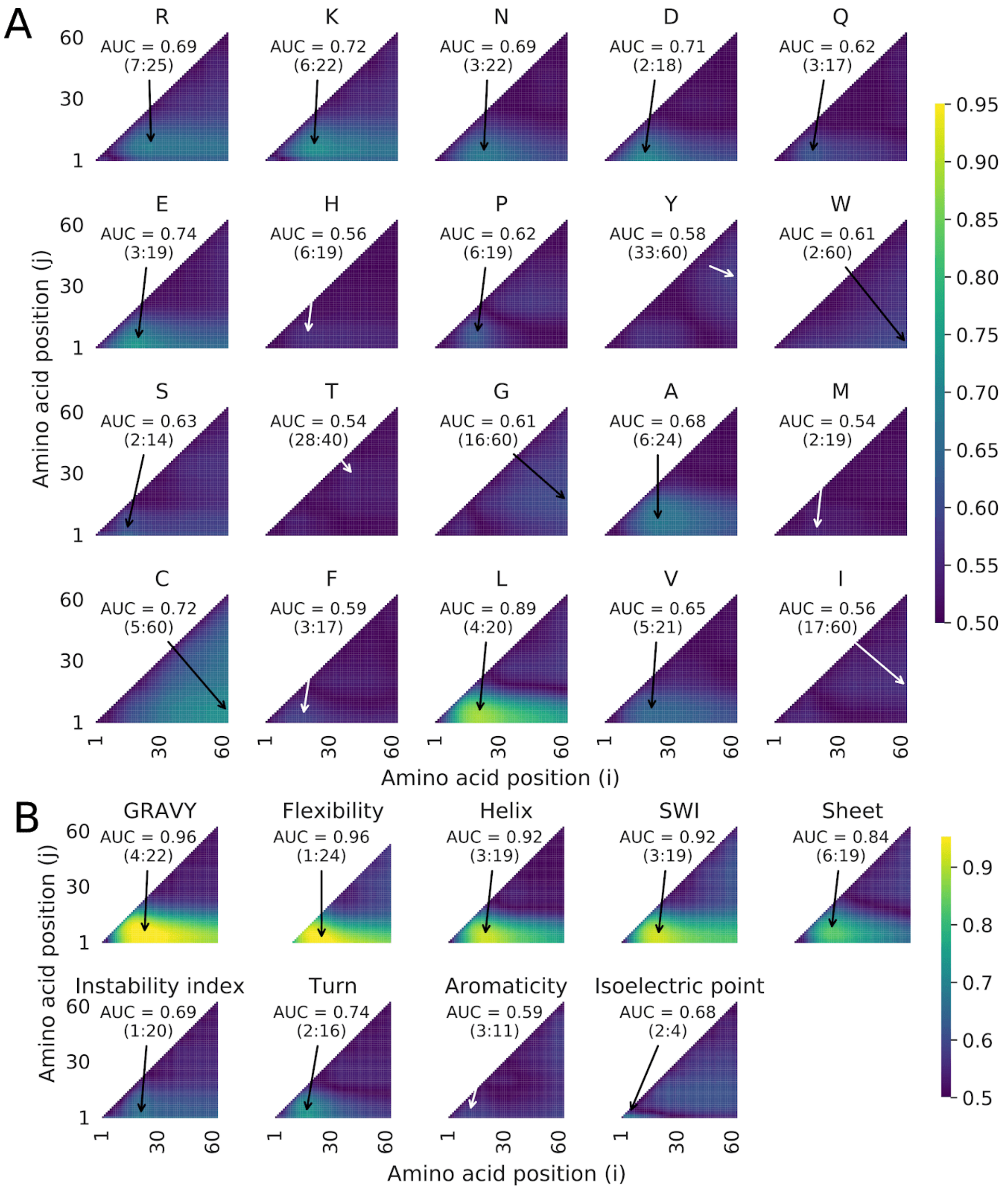


**Fig S2. Leucine (L) composition within the N-terminal region 4:20 shows the highest AUC score in classifying the presence or absence of eukaryotic signal peptides (1,964 and 13,237, respectively).** Amino acid compositions were calculated from positions i to j. **(A)** AUC heatmaps for all residues. **(B)** GRAVY, Flexibility, Helix and SWI are the top four features ranked by AUC scores. AUC, Area Under the Curve; GRAVY, GRAnd average of hydropathicitY; SWI, Solubility-Weighted Index.

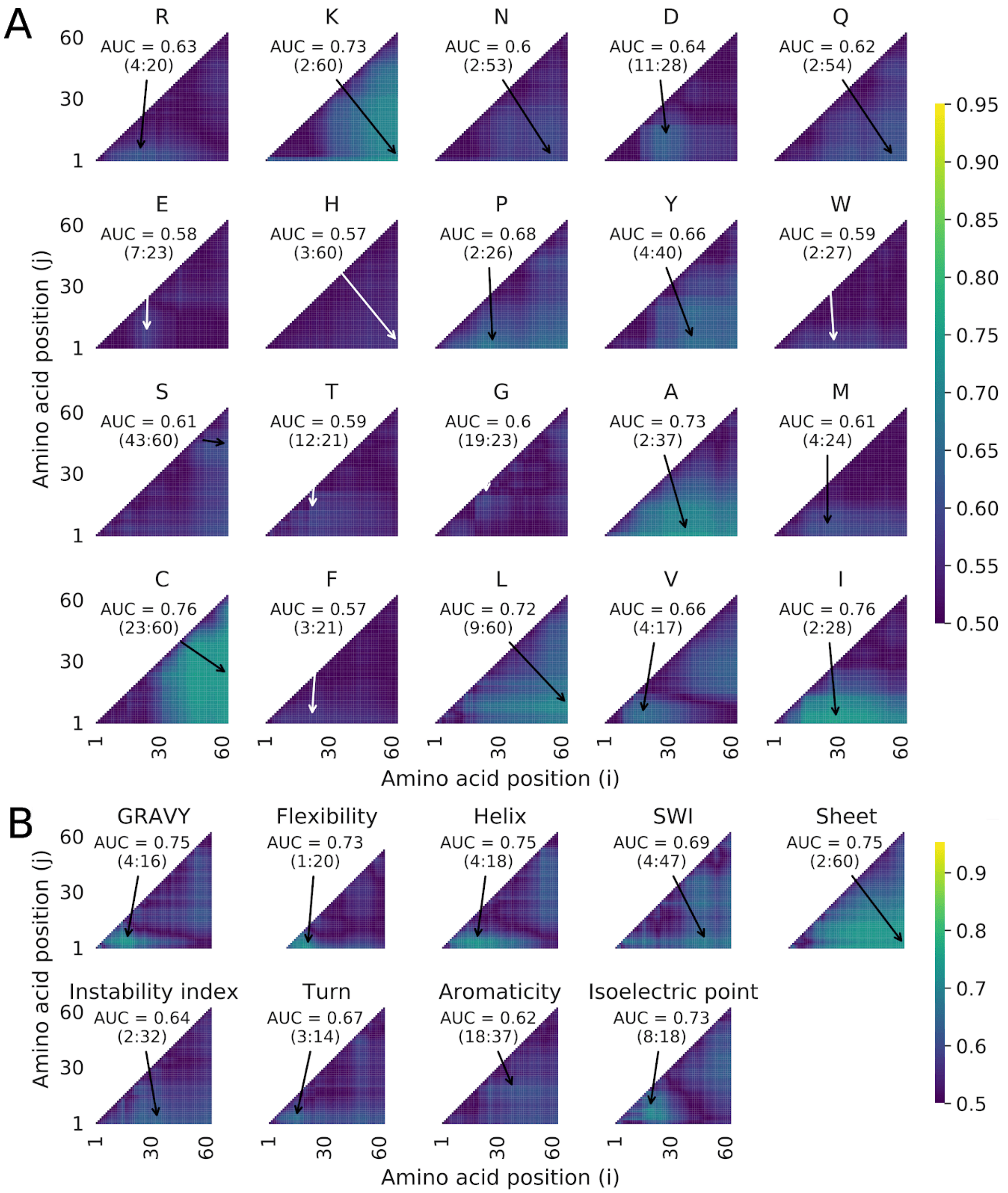


**Fig S3. Isoleucine (I) composition within the N-terminal region 2:28 shows the highest AUC score in classifying toxin and non-toxin SPs (261 and 1,738, respectively).** Amino acid compositions were calculated from positions i to j. **(A)** AUC heatmaps for all residues. Cysteine shows a higher AUC score at the mature region (23:60) as many toxins are cysteine-rich. **(B)** GRAVY, Flexibility, Helix, SWI, Isoelectric point are the top features ranked by AUC scores. Although Sheet has a high AUC score, the region 2:60 extends beyond the normal SP length of around 30 residues. AUC, Area Under the Curve; GRAVY, GRAnd average of hydropathicitY; SWI, Solubility-Weighted Index.

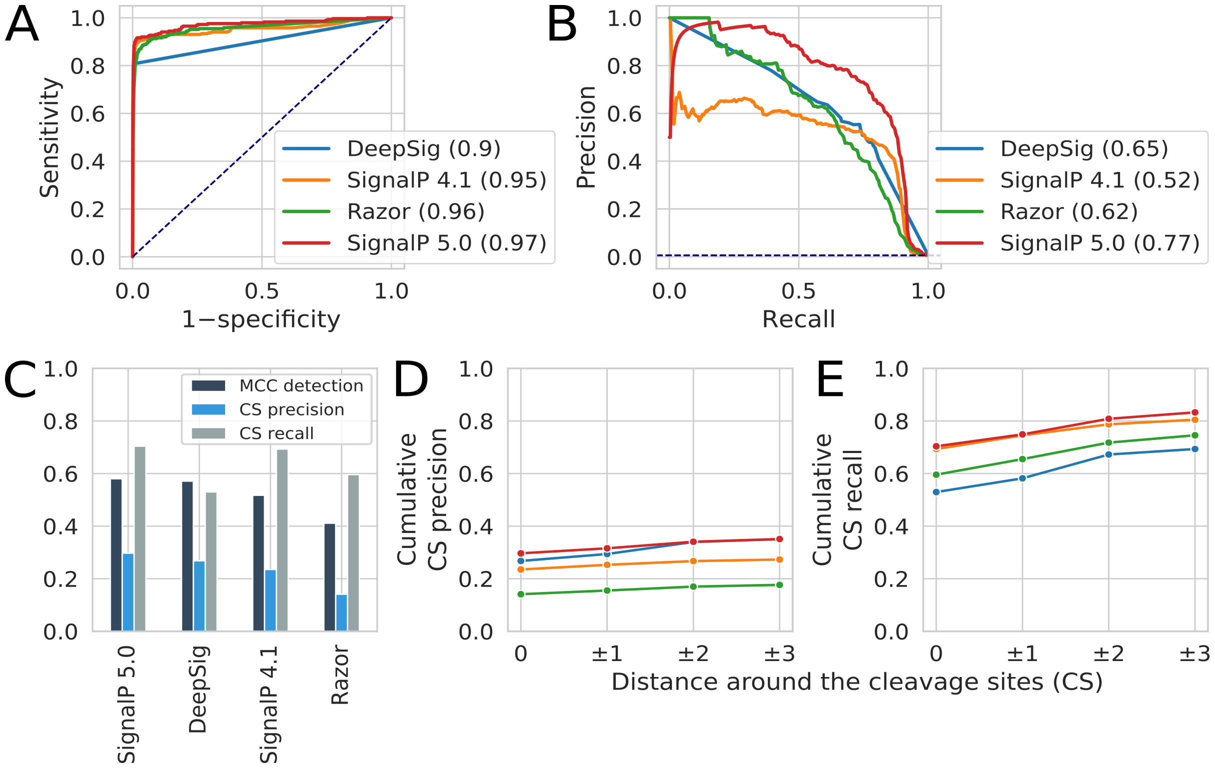


**Fig S4. Performance of Razor and state-of-the-art in predicting eukaryotic SPs using an independent test set (SPs=241, non-SPs=52,055).** Receiver operating characteristic curves **(A)** and precision recall curves **(B)** of the SP prediction tools. Areas under the curves are shown in parentheses. The dotted lines show the performance of a random classifier. **(C)** Matthew's Correlation Coefficients (MCC) of the SP prediction tools. The cleavage site (CS) precisions **(D)** and recalls **(E)** of windows surrounding the cleavage sites are shown. Data are available in Supplementary Tables S3 and S4.

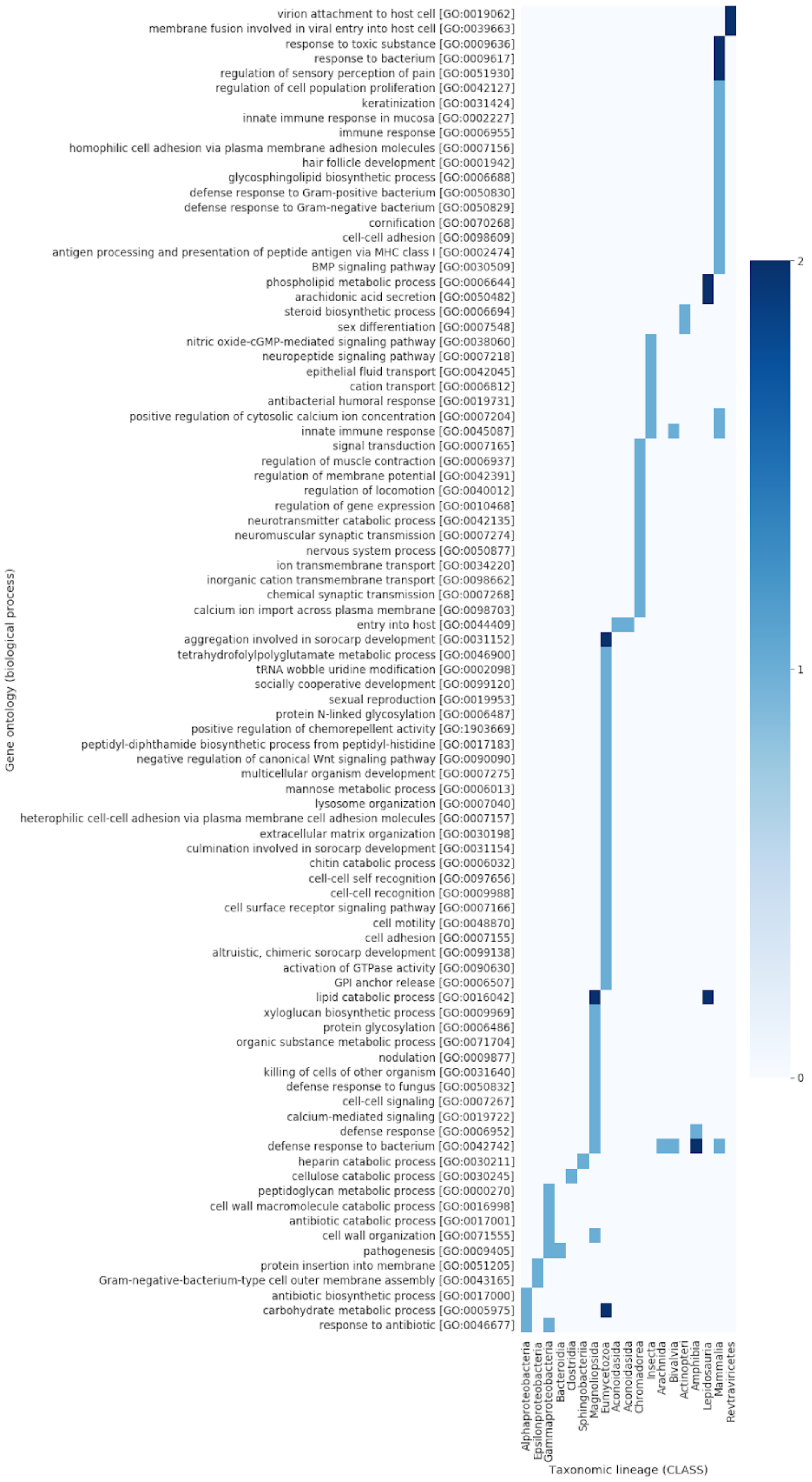


**Fig S5. Gene ontology (GO) annotations (biological process) for the predicted toxin SPs.** A total of 54 out of 100 predicted sequences had GO terms. The scale bar indicates the frequencies of GO terms for the predicted sequences.

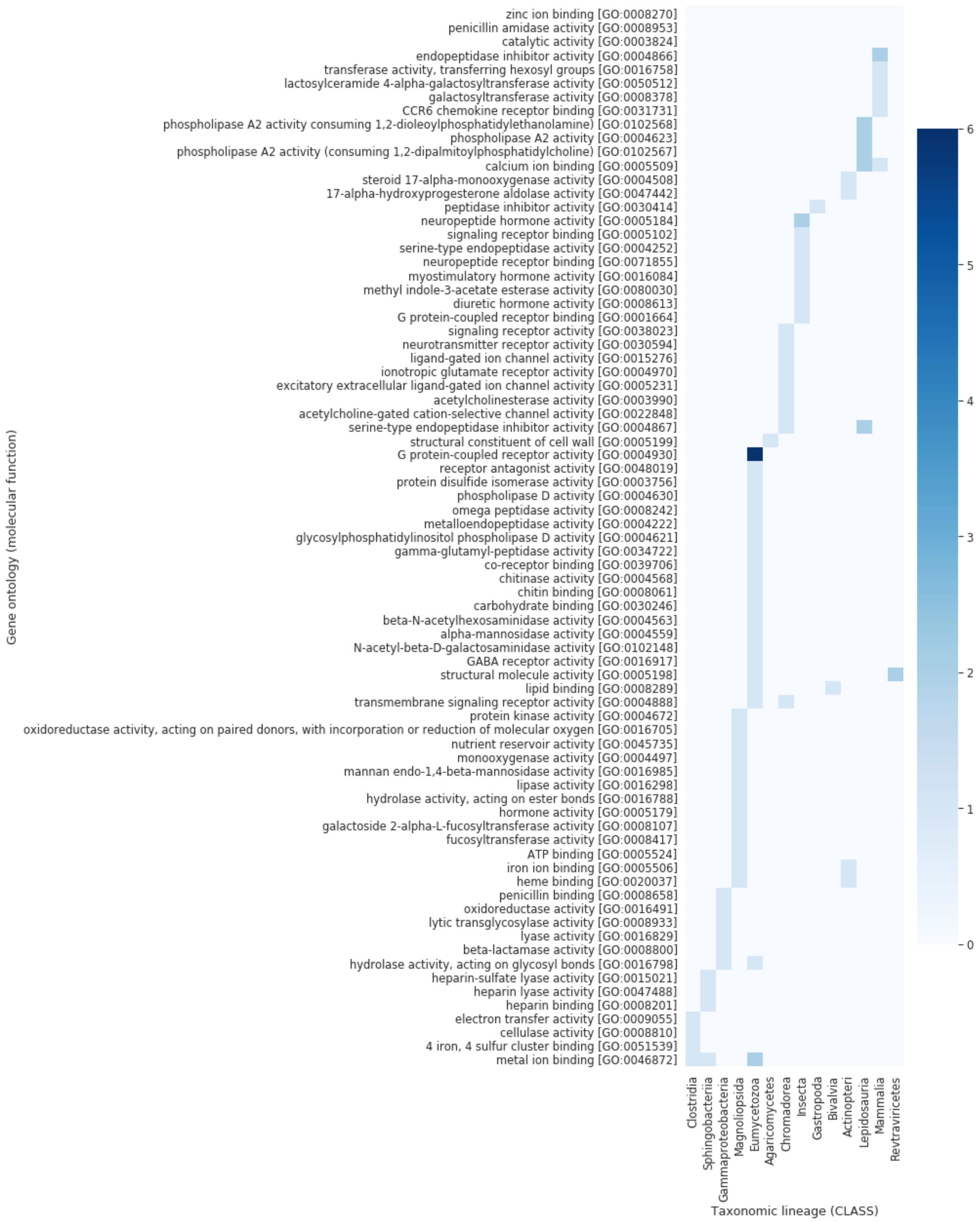


**Fig S6. GO annotations (molecular function) for the predicted toxin SPs.** A total of 59 out of 100 predicted sequences had GO terms. The scale bar indicates the frequencies of GO terms for the predicted sequences.

**Table S1.** Datasets used in this study.

| | **SPs** | **Non-SPs** |
|---|---|---|
| Eukaryotic[a] | 1,694 | 13,237 |
| Toxin[a] | 261 | |
| Independent test set | 214<br>(47 Toxin SPs, 194 Non-toxin SPs) | 52,249 |

[a]Sequences were retrieved from the SignalP 5.0 training set and the animal toxin annotation program of UniProt and were clustered at 70% identity using CD-HIT.

**Table S2.** Feature selection for building the toxin classifier using five-fold cross-validations. Boldface denotes the maximum MCC score.

| **N-terminal lengths** | **MCC scores** | **Features** |
|---|---|---|
| 15 | 0.727 | Hydrophobicity, SWI, Flexibility, Helix, Turn |
| 16 | 0.731 | Hydrophobicity, SWI, Flexibility, Turn, Isoelectric point |
| 17 | 0.716 | Hydrophobicity, SWI, Helix, Turn |
| 18 | 0.726 | Hydrophobicity, SWI, Isoelectric Point |
| 19 | 0.727 | Hydrophobicity, SWI, Turn, Isoelectric Point |
| 20 | 0.727 | SWI, Flexibility, Helix, Turn |
| 21 | 0.718 | Hydrophobicity, SWI, Flexibility |
| 22 | 0.717 | Hydrophobicity, SWI, Isoelectric Point |
| **23** | **0.741** | **Hydrophobicity, SWI, Flexibility, Turn** |
| 24 | 0.715 | Hydrophobicity, SWI |
| 25 | 0.718 | Hydrophobicity, Flexibility, Turn |
| 26 | 0.701 | Hydrophobicity, SWI, Helix, Isoelectric Point |
| 27 | 0.716 | Hydrophobicity, SWI, Flexibility, Isoelectric Point |
| 28 | 0.712 | Hydrophobicity, SWI, Turn |

**Table S3.** Benchmarking of eukaryotic SP prediction using an independent test set (toxin SPs=287, Non-SPs=52,055). Boldface denotes the maximum MCC score.

| | SignalP 5.0 | DeepSig | SignalP 4.1 | Razor |
|---|---|---|---|---|
| MCC | **0.571** | 0.537 | 0.511 | 0.405 |

**Table S4.** Benchmarking of the cleavage site prediction for eukaryotic SPs using an independent test set (SPs=287, Non-SPs=52,055). Boldface denotes the highest scores.

| Tools | Distance around the cleavage sites | | | |
|---|---|---|---|---|
| | 0 | ±1 | ±2 | ±3 |
| Cleavage site precisions | | | | |
| SignalP 5.0 | **0.287** | **0.306** | **0.330** | **0.340** |
| SignalP 4.1 | 0.229 | 0.247 | 0.260 | 0.266 |
| Razor | 0.136 | 0.150 | 0.164 | 0.171 |
| DeepSig | 0.237 | 0.261 | 0.301 | 0.310 |

| Cleavage site recalls | | | | |
|---|---|---|---|---|
| SignalP 5.0 | **0.704** | **0.749** | **0.808** | **0.833** |
| SignalP 4.1 | 0.693 | 0.746 | 0.787 | 0.805 |
| DeepSig | 0.530 | 0.582 | 0.672 | 0.693 |
| Razor | 0.596 | 0.655 | 0.718 | 0.746 |

**Table S5.** Benchmarking of toxin SP prediction using an independent test set (toxin SPs=47, Non-toxin SPs=52,055). Boldface denotes the maximum MCC score.

| | SignalP 5.0 | DeepSig | SignalP 4.1 | Razor |
|---|---|---|---|---|
| MCC | 0.301 | 0.300 | 0.260 | **0.611** |

**Table S6.** Benchmarking of the cleavage site prediction for toxin SPs using an independent test set (toxin SPs=47, Non-toxin SPs=52,055). Boldface denotes the highest scores.

| Tools | Distance around the cleavage sites | | |
|---|---|---|---|
| | 0 | ±1 | ±2 |
| Cleavage site precisions | | | |
| SignalP 5.0 | 0.094 | N/A | N/A |
| SignalP 4.1 | 0.065 | 0.068 | 0.070 |
| Razor | **0.355** | **0.373** | **0.382** |
| DeepSig | 0.073 | 0.077 | 0.097 |
| Cleavage site recalls | | | |
| SignalP 5.0 | **0.979** | N/A | N/A |
| SignalP 4.1 | 0.915 | 0.957 | 0.979 |
| DeepSig | 0.702 | 0.745 | 0.936 |
| Razor | 0.830 | 0.872 | 0.894 |
